# fimpera: drastic improvement of Approximate Membership Query data-structures with counts

**DOI:** 10.1101/2022.06.27.497694

**Authors:** Lucas Robidou, Pierre Peterlongo

## Abstract

**Motivation:** High throughput sequencing technologies generate massive amounts of biological sequence datasets as costs fall. One of the current algorithmic challenges for exploiting these data on a global scale consists in providing efficient query engines on these petabyte-scale datasets. Most methods indexing those datasets rely on indexing words of fixed length *k*, called *k*-mers. Many applications, such as metagenomics, require the abundance of indexed *k*-mers as well as their simple presence or absence, but no method scales up to petabyte-scaled datasets. This deficiency is primarily because storing abundance requires explicit storage of the *k*-mers in order to associate them with their counts. Using counting Approximate Membership Queries (cAMQ) data structures, such as counting Bloom filters, provides a way to index large amounts of *k*-mers with their abundance, but at the expense of a sensible false positive rate.

**Results:** We propose a novel algorithm, called `fimpera`, that enables the improvement of any cAMQ performance. Applied to counting Bloom filters, our proposed algorithm reduces the false positive rate by two orders of magnitude and it improves the precision of the reported abundances. Alternatively, fimpera allows for the reduction of the size of a counting Bloom filter by two orders of magnitude while maintaining the same precision. `fimpera` does not introduce any memory overhead and may even reduces the query time.

**Availability:** https://github.com/lrobidou/fimpera

**Supplementary information:** Supplementary data are available at *Bioinformatics* online.

## 1 Introduction

Public data banks providing sequencing data or assembled genome sequences are growing at an exponential rate [7], and faster than computational power. Searching for a sequence of interest among datasets is a fundamental need that enables for instance a better understanding of genetic changes in tumors, offering precious information about the diagnosis and treatment of cancer [25], or enabling to study at a large scale the distribution and adaptation of life in oceans [24]. However, no method scales to the dozens of petabytes of data already available today. Thus, new computational methods are required to perform a search against datasets.

Querying datasets can be done precisely by aligning genome sequences (e.g. using Blast-like [2] algorithms). However aligning sequences is computational-resources intensive and cannot be applied efficiently when datasets are raw sequencing data. Thus, queries on large-scale datasets are usually done through *k*-mers presence/absence: each dataset is represented by its set of *k*-mers, and a query is represented by its sequence of *k*-mers. The ratio of the *k*-mers in the query and in the dataset over all the *k*-mers in the query approximates the similarity between the query and the dataset. The challenge is to index hundreds of billions of distinct *k*-mers across thousands of datasets. Methodological developments have thus been made to index every *k*-mer of a dataset. Some methods use Approximate Membership Query data structures (AMQ), e.g. Bloom filters, to store the presence/absence of *k*-mers, as for instance SBT [23] or HowDeSBT [10]; see [13] and [6] for a survey of the approaches. However, very few methods tackle the issue of recording the abundance of the indexed *k*-mers. The information about the abundance is however crucial for many biological applications such as in transcriptomics, metagenomics, or metatranscriptomics analyses. We can distinguish three strategies for indexing *k*-mer abundances:

- Explicitly storing couples (*k*-mer, abundances). This may be done simply using hash tables with *k*-mers as keys and their abundances as values. However, this approach cannot scale to dozens or billions of distinct *k*-mers. More compact solutions reduce the *k*-mer set size using assemblies such as compact de Bruijn graph representation (see [4] or [1]) or a spectrum-preserving string set (SPSS) [19]. However, these approaches still require the explicit association of each represented *k*-mer to its set of abundances in each indexed dataset. Additionally, computing the de Bruijn graph or the SPSS of a set of reads requires intensive computational resources. As such, while being effective on e.g. full genomes, it becomes inefficient when representing highly complex and diverse datasets such as metagenomic seawater for instance. Note that once the set of *k*-mer is computed, the memory cost of adding counts is not the bottleneck (see [16], in which the storage of abundances requires only a fraction of the total memory usage). Moreover, counts can be stored as an approximation (see [21]) or exactly, as described below.
- Using a minimal perfect hash function (MPHF) such as [12], or more recently [18]. MPHFs enable the association of an indexed key to a specific and unique value. They provide an efficient way to associate a key to any piece of information. In our context, a *k*-mer can be associated with its abundance in various datasets. Not talking about their construction computation time, these MPHFs do not enable the detection of whether a queried *k*-mer belongs to the original indexed set. Hence, they provide erroneous information for any non-indexed *k*-mer, limiting their usage in this context. To circumvent this problem, certain MHPFs rely on an explicit representation of the indexed *k*-mer set and hence also fall into the previous strategy. This is for instance the case of Blight [14] and SSHash [17]. However, for the same reasons as previously mentioned, these approaches cannot be applied to highly complex and diverse datasets.
- Use an AMQ, adding the abundance information instead of only the presence/absence of each *k*-mer, in this case, we call this AMQ a “counting AMQ”. Using a counting AMQ, the count information cannot be stored using a distinct structure in which the redundant count information between *k*-mers could be compressed, as proposed in [22]. Instead, in a counting AMQ, the abundance of each stored *k*-mer is explicitly represented and thus is not space-efficient. Adding abundance information in an AMQ at fixed memory usage increases its false positive rate. For example, `BIGSI` [5] relies on Bloom filters with a high false positive rate, e.g. 25% false positive rate per *k*-mer query. At constant memory usage, adding the information of abundance using e.g. five bits per cell would yield an extremely high false positive rate that could reach up to 70%, which is not tolerable.

In this paper, we propose a wrapper to improve any existing counting AMQ. The method we introduce is called `fimpera`. It generalizes a previous contribution called `findere` [20]. In short, `fimpera` splits every *k*-mer into *s*-mers (with *k* ≥ *s >* 0) and then associates the abundance of a *k*-mer with its constituent *s*-mers in a counting AMQ. This allows us to retrieve the abundance of a *k*-mer at query time via its *s*-mers count. We show that, when compared to the original counting AMQ indexing of kmers, `fimpera` improves abundance correctness while lowering the false positive rate by an order of magnitude without generating false-negative calls or underestimating the abundance of a kmer and without requiring additional time for query execution. Alternatively, for a fixed false positive rate value, the `fimpera` strategy allows reducing the size of the `cAMQ` by two orders of magnitude.

The `fimpera` algorithm can be used on top of any query made using any kind of counting AMQ. However, our implementation and tests are proposed on top of a counting Bloom Filter (`cBF`) that uses a unique hash function. This choice is motivated by the fact that `cBFs` are the simplest and are widespread data structures for dealing with billions of elements, and by the fact that state-of-the-art indexing tools based on counting AMQ (Cobs [3], HowDeSBT [10]) impose the usage of a unique hash function.

Additionally, the `fimpera` algorithmic needs led us to propose a novel algorithm for computing the sliding window minima (resp. maxima): the minimal (resp. maximums) values of all sub-arrays of a fixed size over an array of *x* values. This algorithm runs in 𝒪 (*x*) time and requires no dynamic memory allocation. This contribution may be useful outside the `fimpera` context. Its novelty is that being in place, it uses no additional memory, while other approaches use memory that is linear with the size of the intervals. This makes it the fastest known algorithm to perform this task. It is available at https://github.com/lrobidou/sliding-minimum-windows along with a benchmark comparing it to other solutions.

The `fimpera` contribution is publicly available at https://github.com/lrobidou/fimpera.

## 2 Methods

### 2.1 Background

A *k*-mer is a word of length *k* over an alphabet Σ. Given a sequence *S*, |*S*| denotes the length of *S*. In the current framework, we consider a dataset to be composed of a multiset of sequences. Given a dataset *D*, 𝒟_*k*_ denotes the multiset of *k*-mers extracted from *D*.

The abundance of a *k*-mer *d* (the number of times *d* appears) in 𝒟_*k*_ is represented by *abundance*(𝒟_*k*_, *d*). We consider that a *k*-mer is “present” in 𝒟 if *abundance*(𝒟_*k*_, *d*) *>* 0, else (*abundance*(𝒟_*k*_, *d*) = 0) the *k*-mer is “absent”.

A counting AMQ data structure represents a multiset of elements 𝒟_*k*_. It can be queried with any element *d*; the query’s response on a counting AMQ, denoted by *n*, is always either correct or overestimated, i.e. *n* ≥ *abundance*(𝒟_*k*_, *d*). If *n* = *abundance*(𝒟_*k*_, *d*), the counting AMQ reports the correct abundance, otherwise it reports an overestimation. Note that underestimation is not possible.

In particular, if *abundance*(𝒟_*k*_, *d*) = 0 and *n >* 0, then *d* is found in the counting AMQ even if it is absent from 𝒟. This particular case is a false positive call. The false positive rate of a counting AMQ, denoted by *FPR*_*cAMQ*_, is defined by 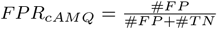 with #*FP* and #*T N* denoting respectively the number of false positive calls and the number of true negative calls (*n* = 0). *FPR*_*cAMQ*_ depends on the used counting AMQ strategy and on the amount of space used by this counting AMQ.

There exist several models and implementations of counting AMQ. The simplest being the counting Bloom Filter [9] (`cBF` for short) using a unique hash function, of which a toy example is given in Fig. 1. In a `cBF`, each element is hashed to get a position in a bit vector, at which position its abundance is stored. This requires a few bits per entry for storing this abundance. Collisions are allowed: should a collision occur, the abundance stored is the maximum of the colliding elements. This leads to a non-null probability of overestimation. A `cBF` is a generalization of Bloom filters, in which each element is allocated only one bit, recording its presence/absence. Hence, a `cBF` requires more memory than a Bloom filter to achieve the same false positive rate. Equivalently, a `cBF` has a higher false positive rate than a simple Bloom filter with constant memory.

**Fig. 1.**
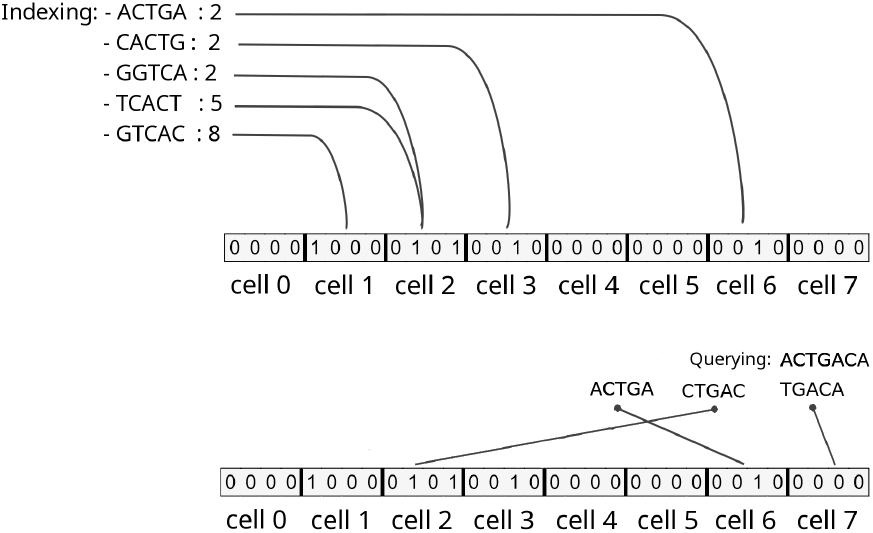
Toy example of a counting Bloom Filter using a unique hash function, with *k*-mers of length *k* = 5, and *b* = 4 bits per element. Top, indexing: each indexed element (*k*-mer) is hashed to a slot, and its abundance is stored in that slot. Bottom, querying: queried elements are hashed, and the value of the corresponding slot is returned. Three situations may occur: 1/true positive (e.g. “ACTGA”), 2/false positive (e.g. “CTGAC”) or 3/true negative (“TGACA”). A true positive is always either correct or overestimated (in case of a hash collision between two elements with different abundances).

It should be noted that other `cAMQ` exist, such as the counting quotient filters [15]. Despite the fact that the method we propose is applicable to any `cAMQ`, the description presented here, as well as the associated implemented tool, are based on the counting Bloom Filter using a unique hash function.

### 2.2 Overview of fimpera

#### 2.2.1 Core principle

`fimpera`’s objectives are to reduce *FPR*_*cAMQ*_ and to improve the precision of the reported abundance of true positive calls. This is achieved using a method based on splitting *k*-mers into smaller words called *s*-mers. The *s*-mers are the elements indexed in the counting AMQ at indexation time. At query time, a *k*-mer is reported as found if and only if every one of its *s*-mers is found in the counting AMQ. Thus, a false positive (resp. overestimation) on a *k*-mer requires *k − s +*1 *>* 1 false positives (resp. overestimations) on its *s*-mers.

Alternatively, for a fixed *FPR*_*cAMQ*_, the `fimpera` strategy reduces the memory needed by the `cAMQ` up to several orders of magnitudes.

Splitting *k*-mers into *z +* 1 *s*-mers and associating each *s*-mer with a hash value as done in `fimpera` may be seen as a similar strategy as using *H* = *z +* 1 hash functions per *k*-mer with a bloom filter. There are three obvious observations to be made here:

- Canonical indexing methods such as HowDeSBT [10] or Cobs [3] for instance rely on bloom filters using a unique hash function. These state-of-the-art approaches can thus directly benefit from our proposal, with no modification.
- Indexing and query time grow linearly with the number of independent hash functions used, while `fimpera` takes advantage of the *s*-mers shared between consecutive *k*-mers. Even if it is not restricted to this framework, `fimpera` is meant for querying streams of *k*-mers. In this situation, when streaming all successive *k*-mers from a sequence, a *k*-mer shares *z s*-mers with its predecessor. Only one unique new *s*-mer has to be inserted or queried for all *k*-mers except the first one. Thus, with `fimpera`, the computation time does not grow with the *z* value. Notably, as shown in Table 2, it even slightly decreases at query time, as once a *s*-mer is detected as absent, all *k*-mers that span this *s*-mer can be skipped.
- Regarding the effect on memory, in a Bloom filter with *H >* 1 distinct hash functions, the *H* value is limited to an optimal as using too many hash functions saturates the filter load factor, leading either to an increase of the false positive rate or to the need of increasing the filter size. This is not the case when using the `fimpera` approach.

#### 2.2.2 Indexation overview

At indexation time, `fimpera` takes as input a file of counted *k*-mers, as provided for instance by KMC [11]. `fimpera` splits each *k*-mer into its *k* − *s* + 1 constituent *s*-mers (*k* ≥ *s*). Each *s*-mer is then stored in a `cAMQ` along with its s-abundance, denoted as *s*_*ab*_. The *s*_*ab*_ of a *s*-mer is formally defined as the maximum of the abundance of the *k*-mers containing this *s*-mer. We explain the choice on relying on *s*_*ab*_ instead of the abundance of *s*-mers and describe its outcomes in the Supplementary Materials, Section 3.3. Using both the *s*_*ab*_ and the abundance of *s*-mers is supported in the implementation.

In the following, we set *z* = *k − s*, hence *z* ≥ 0.

#### 2.2.3 Query overview

For each queried sequence *S*, `fimpera` extracts its sequence of *s*-mers, which is then queried against the `cAMQ`, and the abundance of any *k*-mer of *S* is computed as the minimum of *s*_*ab*_ of its *s*-mers. By default, `fimpera` prints each input sequence along with the abundance of every of its consecutive *k*-mer. In practice, the input query file is a fasta or a fastq file, possibly gzipped.

#### 2.2.4 False positive calls

Let’s consider a *k*-mer *d* with an abundance of 0 and each of its *s*-mer has an *s*_*ab*_ of 0 as well. With `fimpera`, wrongly reporting *d* as present requires that *every s*-mer of that *k*-mer are wrongly found as present in the counting AMQ. The probability of such an event is approximately *F* (*PR*_*cAMQ*_)^*z*+1^, leading to a dramatic decrease in the occurrences of false positive calls with respect to *z*. For instance, with *z*=3 (which is a recommended and the default value) and a counting AMQ having a false positive rate of 25%, the false positive rate with `fimpera` for that setting is ≈ 0.4%.

The `fimpera` approach may generate a novel kind of false positive. A queried *k*-mer, absent from the indexed dataset, may be composed of *s*-mers existing in this indexed set. Querying such a *k*-mer with `fimpera` returns a non-zero abundance, so generating a false positive, which we call a “construction false positive”. These false positives are created by `fimpera`, *independently of the underlying* `cAMQ`. This event is non-null but it is in practice negligible when using usual *k* and *z* values, as shown in the results.

#### 2.2.5 Overestimations

To overestimate the abundance of a queried *k*-mer with `fimpera`, overestimations are required to happen on the abundance of every *s*-mer of that *k*-mer that has the minimal *s*_*ab*_. The more *s*-mer per *k*-mer, the more *s*-mer abundance overestimations need to happen to overestimate a *k*-mer abundance. *s*-mer abundance overestimations can come from two sources:

- a collision occurs in the counting Bloom Filter, leading to the overestimation of the less abundant colliding *s*-mer; and/or:
- a *s*-mer is shared among two different *k*-mers having different abundances. This overestimates the abundance of this *s*-mer from the least abundant *k*-mer. This happens no matter the false positive rate of the counting Bloom Filter. We call those overestimations “construction overestimation”. This new kind of overestimation is specific to `fimpera`.

Observe a case of interest: consider two *k*-mers *d*_0_ and *d*_1_ overlapping over *k −* 1 characters. If *d*_1_ has an abundance greater than *d*_0_, then the correct abundance of *d*_0_ is retrievable through a unique *s*-mer (the unique *s*-mer of *d*_0_ that does not appear in the *k*-mer *d*_1_). In such case, *d*_0_ is more likely to be overestimated than *d*_1_.

Consequently, `fimpera`’s overestimations are not uniformly distributed random events. Overestimations are more likely to occur close to a change in abundance along queried sequences than in a random *k*-mer. In such cases, those overestimations are limited to the abundance of their neighbor *k*-mers, mitigating their impact. Indeed, in the result section (Section 3), we show that the erroneous abundance calls are closer to the ground truth with `fimpera` compared to those obtained with the original `cBF`.

We now describe in more detail both the indexing step and the querying step of `fimpera`, as well as one optimization, allowing a query time independent from the *z* value.

Another optimization enables `fimpera` to perform queries slightly faster than using the original underlying `cAMQ`. This optimization is described in Supplementary Materials, Section 2.2.

### 2.3 Querying with fimpera

fimpera’s query consists in querying all consecutive, overlapping *k*-mers from a sequence of size greater than or equal to *k* through their constituent *s*-mers. `fimpera`’s query is a two-step process:

- for every position in the query except the last *s −* 1 ones, *s*-mers starting at these positions are queried in the counting AMQ and are stored in an array of integers *s*_*ab*_;
- the abundance of any *k*-mer starting position *p* is the minimum value of the sub-array of length (*z +*1) starting at the position *p*: *s*_*ab*_[*p*; *p + z*].

A non-optimized version of the `fimpera`’s query algorithm is shown in Supplementary Materials, Section 1.

The first step is improved by avoiding recomputing the minimal value of windows of length *z +*1 starting at each position *p* of an array. This optimization is described in the following section.

### 2.4 Optimisation: sliding window minimum algorithm

The problem, independent of `fimpera`, is as follows: given a vector of values (integers or floats) *v* and an integer *W* (denoting the size of a sliding window, with *W* ≤ |*v*|), give an array *r* such that ∀*i* ∈ [0, |*v*| − *W*], *r*[*i*] = *min*(*v*[*i*], *v*[*i +* 1], …, *v*[*i + W −* 1]). This is a particular case of a more general problem, the “range minimum query” (RMQ). Given a vector *v* of element drawn from a totally ordered set and two integers *i, j* (0 ≤ *i < j <* |*v*|), the RMQ consists in finding *min*(*v*[*i*], *v*[*i* + 1], …, *v*[*j −* 1]. Answering RMQ generally relies on some pre-computation beforehand (e.g. pre-computing the whole set of possible queries, but less resource-intensive solutions can be found, such as [8]).

The sliding window minimum problem studied here is a particular case of an RMQ (*j − i* is fixed and queries consist in every window of that size). Thus, its solutions do not require taking into account other window sizes, effectively allowing to skip pre-computation. The naive approach to solving the sliding minimum problem is to simply search for the minimal value in each window. This algorithm is in 𝒪 (*W* × |*v*|) time. Some straightforward faster solutions can be built on top of dynamic heaps. Here, we propose a solution (named “fixed window”) that does not rely on any heap allocation (which is slow for most systems). This solution without any dynamic memory allocation is an order of magnitude faster than other 𝒪(|*v*|) solutions as shown Section 3.1 in the Supplementary Materials.

The main idea of the proposed “fixed window” approach is to split the input vector of values in fixed, non-overlapping windows of size *W*. Then, for each so-called “fixed window”, compute two vectors:

- *min*_*L*__*j*: *min*_*L*__*j*[*i*% *W*] contains the minimum value encountered in the *j*-th fixed window up to the position *i*;(*i* ∈ [*i* × *W* × *j*; *i* × *W* × (*j +* 1) − 1]
- *min*_*R*__*j*: *min*_*R*__*j*[*i*%*W*] contains the minimum value from the position *i* up to the end of the *j*-th fixed window.

All *min*_*L*__*j* and *min*_*R*__*j* vectors are then concatenated into two vectors (*min*_*L*_ and *min*_*R*_). The minimum of a *sliding* window starting at position *i*, denoted by *r*[*i*], is thereupon the minimum between:

- *min*_*L*_[*i + W −* 1] (the minimum of the left part of the next fixed window)
- *min*_*R*_[*i*] (the minimum of the right part of the current fixed window)

An example is provided in Table 1.

**Table 1.** Computation example of the *r* vector, with a window of size 3. Tables *min*_L_ and *min*_R_ are represented for helping comprehension but are not implicitly created in practice. The *j* row indicates the starting positions of each fixed windows. As an example, the minimal value of the sliding window of size 3 starting position *i* = 1 is *r*[1] = 1 (bold underlined value), being equal to min(*min*_L_[1 + 3 − 1], *min*_R_[1]) = min(1, 3) (underlined values).

|  |  |  |  |  |  |  |  |  |  |  |
| --- | --- | --- | --- | --- | --- | --- | --- | --- | --- | --- |
| $i$ | 0 | 1 | 2 | 3 | 4 | 5 | 6 | 7 | 8 | 9 |
| $v$ | 5 | 3 | 7 | 1 | 4 | 5 | 3 | 2 | 2 | 3 |
| $j$ | 0 | | | 1 | | | 2 | | | 3 |
| $\min_L$ | 5 | 3 | 3 | <u>1</u> | 1 | 1 | 3 | 2 | 2 | 3 |
| $\min_R$ | 3 | <u>3</u> | 7 | 1 | 4 | 5 | 2 | 2 | 2 | 3 |
| $r$ | 3 | **<u>1</u>** | 1 | 1 | 3 | 2 | 2 | 2 | | |

**Table 2.** Influence of the *z* parameter on the quality of the results and on the computation time when indexing and querying 31-mers through their *s*-mers. “constr.” stands for “construction” and “overest.” stands for “overestimation”. Results with *z* = 0 are equivalent to those obtained with the original `cBF` results. The Incorrect abundance calls are computed over true positive calls only.

| $z$ | 0 | 3 | 5 | 7 | 9 | 20 |
| --- | --- | --- | --- | --- | --- | --- |
| (indexed $s$-mer size $s = 31 - z$) | 31 | 28 | 26 | 24 | 22 | 11 |
| False positive rate (%) | 25.00 | 0.56 | 0.08 | 0.02 | 0.03 | 99.70 |
| Of which: construction FP (%) | 0 | 0.49 | 9.64 | 46.32 | 67.26 | 99.99 |
| Incorrect abundance calls (%) | 1.55 | 1.33 | 1.93 | 2.99 | 4.27 | 42.73 |
| Of which: constr. overest. (%) | 0 | 83.02 | 88.65 | 94.24 | 94.34 | 99.85 |
| Query time (s) | 557 | 514 | 498 | 491 | 493 | 528 |

Note that, as described previously, this approach would require allocating memory for two vectors per call. This memory need may appear negligible in theory as those vectors are limited by the query size which is a few hundred to few thousand. However, in practice, allocating memory for these vectors is time-consuming, and may increase significantly the practical running time. We overcame this memory need thanks to these three following tricks. 1/we compute *min*_*L*_[*i*] on the fly (if *i*% *W ≠* 0 then *min*_*L*_[*i*] = *min*(*min*_*L*_[*i −* 1], *v*[*i*]), else *min*_*L*_[*i*] = *v*[*i*]). 2/*min*_*R*_ is computed *directly in the queried vector*. This does not impact the correctness of the algorithm, as *r*[*i*] ≤ *min*_*R*_[*i*] ≤ *v*[*i*]. 3/ the response (minimal value per sliding window) can be stored directly in the input queried vector as well. At the price of modifying the input vector, this allows the algorithm to be run in 𝒪 (*size*_*query*) time while avoiding any time-consuming heap allocation.

A complete description of the optimized solution is provided in Supplementary Materials, Section 2.1, algorithm 2, along with a benchmark Section 3.1, Fig. 1.

This algorithm offers a generic solution for computing the minimal value of a sliding window in constant memory and linear time. Its usefulness is not limited to `fimpera`. As so, we propose an independent implementation https://github.com/lrobidou/sliding-minimum-windows. Note also that it can be straightforwardly modified for computing the maximal value instead of the minimal value of each window.

### 2.5 Implementation of fimpera

An implementation of `fimpera` is available at https://github.com/lrobidou/fimpera. This implementation is specialized for genomic data (i.e. with an alphabet consisting of A, T, C, G) and uses a counting Bloom Filter with a unique hash function as `cAMQ`. A template mechanism allows the use of any other `cAMQ` provided by the user. Queries consist of fasta or fastq files (gzipped or not), and an option is provided to index and query canonical *k*-mers only, i.e. the lexicographic minimum between each *k*-mer and its reverse complements. Indexing options include the *k* and *z* values, the size of the filter, and *b*, the number of bits per element used to store its abundance. As *b* has a major impact on the final size of the data structure, it is recommended to use low *b* values (say *b* ≤ 5). This limits the maximal stored abundance value to 2^*b*^ − 1 (to prevent overflow, the abundance is capped at 2^*b*^ *−* 1).

In order to encode large abundance values with few bits per *k*-mer, instead of storing the first possible 2^*b*^ − 1 distinct abundance values, we propose to discretize any abundance value to user-defined interval ranges.

In practice, `fimpera` can use any surjective function

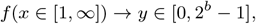

supplied by the user. This can be for instance any intervals. The proposed implementation proposes the usage of *y* = min([log_2_(*x*)], 2^*b*^ − 1), or *y* = min([log10(*x*)], 2^*b*^ − 1).

Changing the default output (e.g. storing results instead of printing them, computing average abundance per sequence, or printing only sequences whose average *k*-mer abundances is above a user-defined threshold) is possible.

## 3 Results

### 3.1 Experimental setup

To the best of our knowledge, no other tool focuses on reducing the false positive rate of existing `cAMQ`, thus we compare `fimpera` results applied on a counting Bloom Filter indexing *s*-mers with the original counting Bloom Filter results indexing *k*-mers using a unique hash function. We propose results on biological marine metagenomic data.

A list of commands for reproducing the results is available here: https://github.com/lrobidou/fimpera/blob/paper/paper_companion/Readme.md along with a step-by-step explanation of the output. Executions were performed on the GenOuest platform on a node with 4×8cores Xeon E5-2660 2,20 GHz with 200 GB of memory.

### 3.2 Metagenomic dataset

We used two fastq files from the TARA ocean metagenomic dataset [24] to show the advantages offered by `fimpera` on metagenomic samples. The index was computed from the 2.38 × 10^8^ distinct 31-mers present at least twice in an arctic station (accession number ERR1726642) and the query sample was the first 3 × 10^6^ reads from a sample in another arctic station (accession number ERR4691696). Canonical *k*-mers were considered for this experiment.

### 3.3 Choice of filters parameters

In this experiment, we apply the `fimpera` approach on top of a counting Bloom Filter designed to have 25% of false positive calls, while using 5 bits per element for storing the abundance of indexed *k*-mers. For indexing 2.38 × 10^8^ 31-mers this structure requires 6.96 × 10^8^ slots (thus 3.48 × 10^9^ bits).

We compare the results of queries made against this raw counting Bloom Filter with the results of queries made using `fimpera` wrapping that same counting Bloom Filter.

Thus, parameters used are as follow: *k* = 31, size of the filter of 3.48×10^9^ bits, as discussed in section 3.3, using *b* = 5 bits per abundance count (thus 2×10^8^ slots), and abundances are stored as their [log_2_]values. We use the default *z* = 3 parameter (unless otherwise stated). As we use *z* = 3, we compare the results of a `cBF` indexing 31-mers, with results of `fimpera` used on a `cBF` with the same sizing, but indexing *s*-mers of size 28 (31-3).

### 3.4 Used metrics

To measure the quality of the `fimpera` results and the `cBF` results, we propose three metrics:

- the false positive rate, which provides the probability that the method returns an abundance call *>* 0 for a *k*-mer absent from the indexed set.
- the proportion of incorrect abundance, that provides the probability that the method returns the incorrect abundance for a *k*-mer actually in the indexed set.
- statistics of responses for incorrect abundances calls, that estimate the reported abundance of *k*-mers whose abundance is incorrectly reported. When comparing any two approaches, we set up a metric that we call “overestimation score”. It is defined as the sum of the square of errors made on the output of each method.The lower the overestimation score is, the fewer errors were made. Errors that are closer to the ground truth are less penalized by this score than errors distant from it.

### 3.5 False positive rate analyses

Results about false positives obtained with the proposed experiment are shown in Fig. 2. Results about the `cBF` simply confirm the setup and show a false positive rate of 25%. When applying `fimpera`, the false positive rate drops to 0.56%. Among all these `fimpera` false positives, 4.8 % are due to the so-called “construction false positives” (see Section 2.2.3), thus representing 0.0027% of the total *k*-mer calls.

**Fig. 2.**
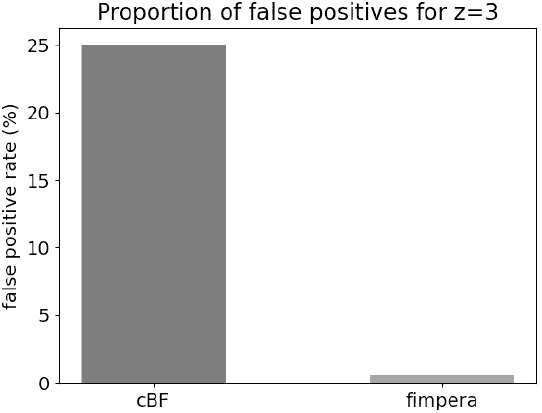
Proportion of false positive calls without `fimpera` (on a classical counting Bloom Filter) and with `fimpera` (*z* = 3), indexing and querying real metagenomic read datasets.

It is important to recall that these comparative results were obtained using the exact same amount of space. Hence the `fimpera` approach enabled to yield about 45 times fewer false positive calls, with no drawback and even saving query time (see Table 2).

#### 3.6 Correctness of the reported abundances

In this section, we focus only on true positive calls. Hence, these results do not concern the 25% false positive calls obtained with the original `cBF`, nor the 0.56 % ones using `fimpera`.

Results comparing the proportion of calls reported with an incorrect abundance among the true positives show that 1.54 % of true-positive calls are overestimated in the `cBF`, while 1.33 % of true-positive calls are overestimated with `fimpera`. Among the `fimpera` calls estimating an incorrect abundance among the true positives, 83 % are due to the so-called “construction overestimation”.

### 3.7 Distribution of errors in overestimated calls

In this section, we focus only on the wrongly estimated calls among true positives.

Results presented Fig. 3 show that, as stated in Section 2.2.5, the erroneous abundance calls are closer to the ground truth with `fimpera` compared to those obtained with the original `cBF`. As seen Fig. 3-bottom, with `fimpera`, almost all (except for a few outliers) overestimations are only one value apart from the correct range (the average difference with the correct abundance range is 1.07). With the original `cBF`, as seen Fig. 3-left, overestimations are more important (1.33 range in average from the ground truth).

**Fig. 3.**
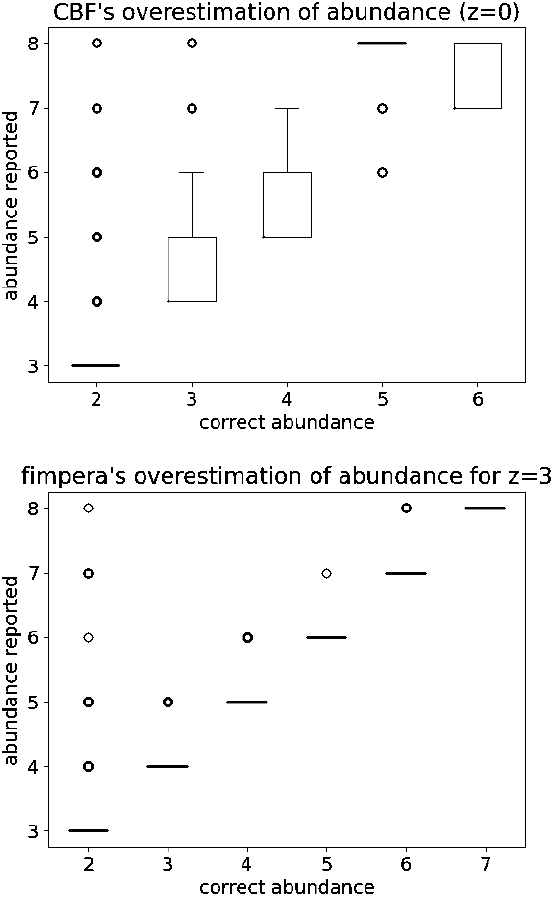
For true-positive calls with an incorrect abundance estimation: reported abundance with respect to the correct abundance. Top: using the original `cBF`, Bottom: using `fimpera`.

### 3.8 Influence of the size of the underlying cAMQ

Because there is a trade-off between space usage and the false positive rate of a `cAMQ`, `fimpera` can either reduce the false positive rate of the `cAMQ` without changing its size or reduce its size without changing its false positive rate. This section focuses on the impact of `fimpera` on this latter strategy.

This trade-off (with and without `fimpera`) is shown Fig. 4 and Fig. 5. To be as close as possible to real-life use-case, we consider in this section the average abundance of *k*-mer on each read, not the abundance of each *k*-mer. For precision requirement, abundances were not stored as their [log_2_] values, but rather as their original values.

**Fig. 4.**
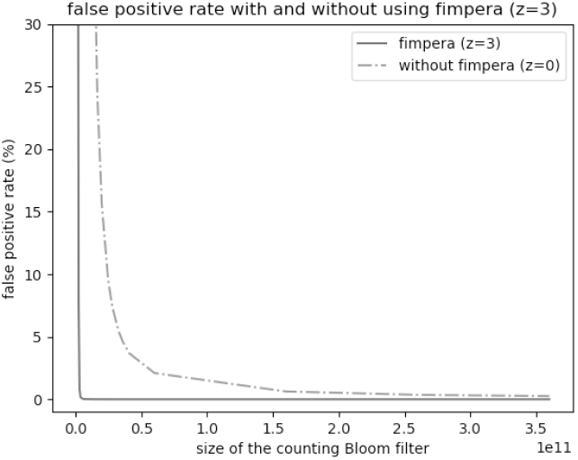
Variation of the false positive rate with respect to the size of the `cBF`, with and without `fimpera` (*z* = 3). As we consider here the average abundance of *k*-mers on each queried read, a false positive means that the average abundance of a *k*-mer on a read is strictly positive while it should have been 0.

**Fig. 5.**
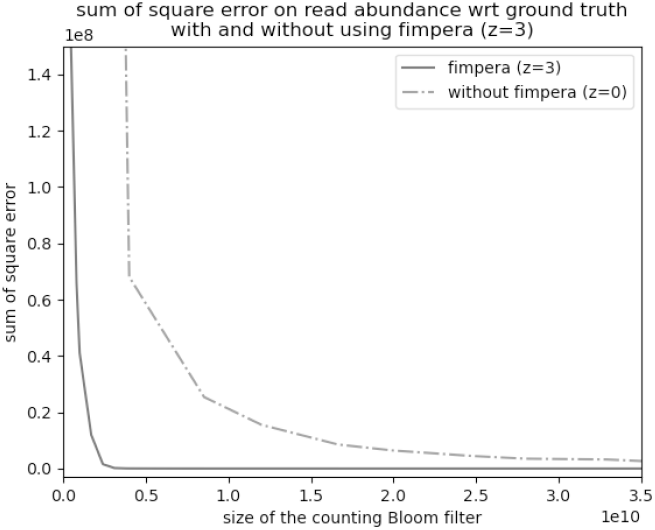
Sum of square error with respect to the size of the `cBF`, with and without `fimpera` (*z* = 3). In this experiment, the output of `fimpera` is the average of each input read.

With `fimpera`, achieving a false positive rate of 2% requires ≈ 2.98 × 10^9^ bits (≈ 3.72 × 10^8^ slots of 8 bits each) (see Fig. 4). Without `fimpera`, achieving the same false positive rate requires ≈ 65.5 × 10^9^ bits (21 times more space). The lower the false positive rate, the greater the space gap between using and not using `fimpera`. Achieving a false positive rate of 0.02% requires about 7.25 × 10^9^ bits with `fimpera`, but 360 × 10^9^ bits were not enough without `fimpera` (i.e. allocating more than 50 times the space budget of `fimpera` would be required). In order to take into account the abundance information, we set up the output of `fimpera` to be the average of *k*-mers’ abundance for each queried read.

Fig. 5 shows the sum of square of overestimations, with and without `fimpera`. For an overestimation score (defined Section 3.4) of 200000, `fimpera` requires ≈ 3.10 × 10^9^ bits (≈ 3.87 × 10^8^ slots of 8 bits each), whereas a `cBF` would requires 330.9 × 10^9^ bits (i.e. 100 times more space).

#### 3.8.1 Impact of z

As shown in Table 2, the false positive rate decreases with respect to *z* and stays low for a wide range of *z* values (at least from *z* = 3 to *z* = 9). When using an extreme *z* value, for instance, *z* = 20, the false positive rate is increased up to almost 100%. With *z* = 20, as we use *k* = 31, the size of the indexed *s*-mers is *s* = 11. When indexing as little as a few hundred million characters, each 11-mer has a great probability to appear by chance in the indexed dataset. Indeed, the probability of such event is 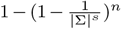 with *n* being the number of characters. For instance, for *s* = 11 and a sequence as little as *n* =20 million characters, the probability of the presence of any 11-mer is *>* 99%. This quasi-random existence of all *s*-mers generates a huge amount of construction false positives, as seen in the last column. This has an effect on the running time, which is close to the query time of *z* = 0, undoubtedly because all *s*-mers queried are positives, annihilating the *s*-mer skipping optimization.

Caveat: in Table 2, the overestimations are reported only for true positive calls. Overall, also taking into account the negative answers, the overestimation rate with `fimpera` would be 0.03 with *z* = 3 for instance.

In the Supplementary Materials, section 3.4, we show the impact of *z* in a different setup: choosing a fixed *s* and increasing *k* with regard to *z*, instead of fixing *k* and decreasing *s* with regard to *z*. This shows another framework for using `fimpera`, with similar conclusions about the quality of the results.

#### 3.8.2 Query time

As mentioned in the Supplementary Materials, section 2.2, the `fimpera` approach does not increase the query execution time. On the contrary, it makes it possible to reduce the running time slightly as *z* increases. See Table 2.

#### 3.8.3 Discussion

A Bloom filter indexing 2.38 × 10^8^ elements using 2.33 × 10^8^ bits and one hash function would lead to a false positive rate of ≈ 6.61%. A counting Bloom Filter indexing the same dataset using the same space with 5 bits per slot (thus 5 times fewer slots) yields a false positive rate of 25%. With `fimpera`, using 5 bits per slot and the same total number of bits, the false positive rate drops to 0.56%. Using the same space as a Bloom filter, `fimpera` allows reducing the false positive rate, while adding count storage and being quicker to query. The only downside may in some cases be the computation of the *s*_*ab*_ from the abundance of counted *k*-mers. However, this can be skipped altogether with a moderate impact on the result (see Supplementary Materials, Section 3.3).

## 4 Conclusion

We presented `fimpera`, a novel computational method for reducing the false positive rate and increasing precision in any counting Approximate Membership Query data structure. This is achieved without requiring any changes to the original data structure, with no memory overhead, and even with a slight improvement in query computation time.

Our results showed that when applied on top of a counting Bloom Filter, `fimpera` enabled to yield about 45 times fewer false positive calls than when querying directly a counting Bloom Filter of identical size. Moreover, using `fimpera`, abundance errors were slightly less frequent on true positive calls, and finally, those abundance errors were on average 1.07 apart from the ground truth with `fimpera` while they are on average 1.33 apart from the ground truth with the original `cBF`.

Independently from parameters of the used `cAMQ`, `fimpera` requires setting up a unique parameter, *z*. Fortunately, results are highly robust with the choice of *z*, unless extreme values are chosen. Future work will include a formal analysis of the theoretical limits on the choice of *z* usage ranges.

We provide a C++ implementation of `fimpera` which enabled us to validate the approach. This implementation can also be used as a stand-alone tool for indexing and querying genomic datasets, and it can be tuned with user-defined parameters and ranges of abundances. The provided GitHub project also proposes all necessary instructions and links to genomic data to reproduce the results.

Finally, of independent interest, we proposed a novel algorithm and its implementation for computing the minimal or maximal values of consecutive windows, sliding on an array of integers or floats. To the best of our knowledge, this is the fastest algorithm to perform this task.

## Acknowledgements

The authors thank Eric Pelletier for his help regarding the usage of the Tara Ocean data sets. We acknowledge the GenOuest bioinformatics core facility (https://www.genouest.org) for providing the computing infrastructure. The work was funded by ANR SeqDigger (ANR-19-CE45-0008).

## Supplementary Materials

These supplementary materials first propose a detailed description of the proposed query algorithms. We start by providing the simple naive query algorithm (Section 1). Then, we present two optimization approaches (Section 2.1 and 2.2). These supplementary materials also propose additional results:

- showing the advantages of the sliding window algorithm we propose 3.1,
- showing a comparison between using `fimpera` on a counting AMQ having a unique hash function, or not using `fimpera` on a counting AMQ with *z* + 1 hash functions (Section 3.2),
- showing the effects of using the abundance of *s*-mers, instead of their s-abundances (Section 3.3),
- showing the effects of choosing the *k* value at query time and not at indexing time (Section 3.4),
- and showing the effects of not grouping values using log functions (Section 3.5).

### 1 Query algorithm: non-optimized version

A non-optimized algorithm of `fimpera`’s query is shown in Algorithm 1. This algorithm takes a queried sequence *q*, a counting AMQ indexing *s*-mers, and parameters *k* and *z*. It returns a vector of integers called *response*, such that: ∀*i* ∈ [0, |*response*| − 1], *response*[*i*] is the abundance of the *k*-mer starting at position *i* in the query, for all *i* in [0, |*q*| − *k* + 1].

#### Algorithm 1

Non optimized `fimpera’s` query

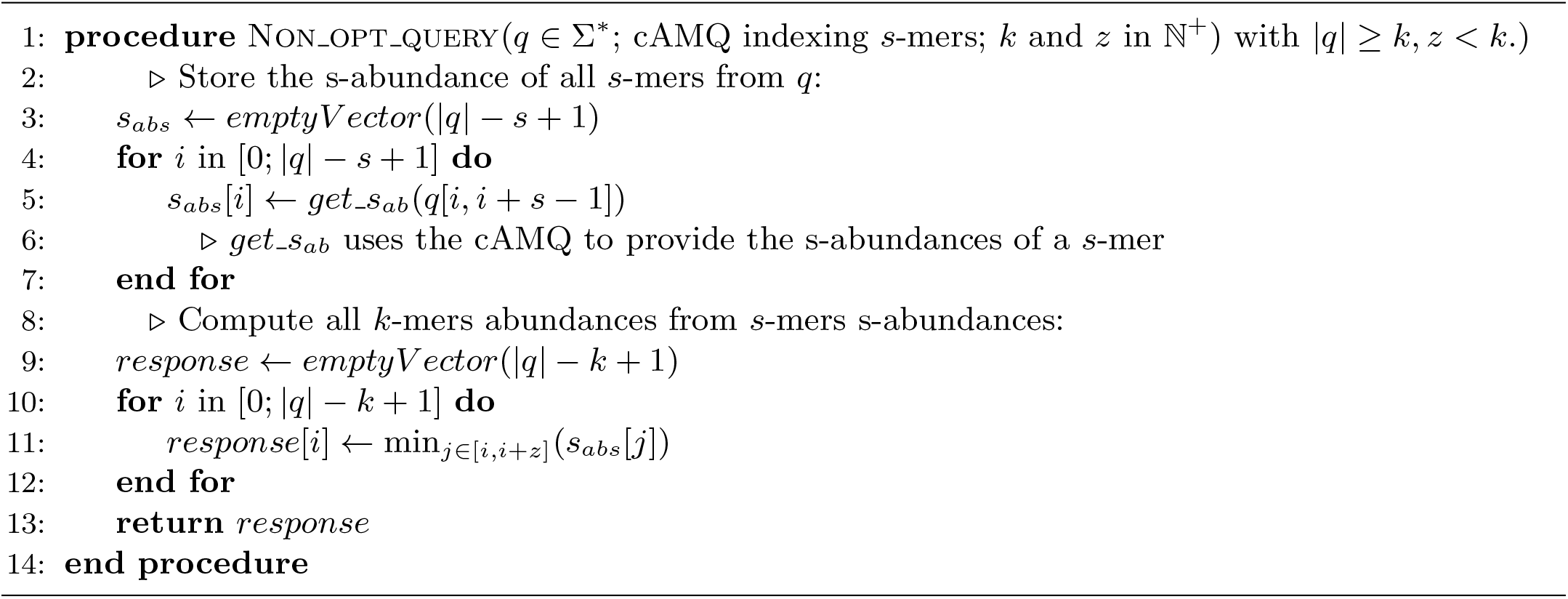

Algorithm 1 is not optimal. Line 11 computes the minimal value of a range of *z* consecutive integers taken from a vector of integers. At each iteration of the *for loop* line 10, this range is shifted by one.

### 2 Optimized query algorithm

We propose two optimizations of the previous algorithm.

- The first one consists in using a minimum sliding window, that provides a way to compute in *O*(1) the *min* from Line 11 of the non-optimized version of the query algorithm. This optimization is explained at a high level in the main text (Section 2.4). Here, Section 2.1 we provide the associated pseudo-code.
- The second optimization consists in exploiting the fact that for any queried *s*-mer detected as absent, all *k*-mers that contain this *s*-mer are also predicted as absent. Section 2.2 shows how to optimize the queries jumping all *k*-mer positions that contain an absent *s*-mer. This section finally proposes the overall optimized query algorithm 3.

#### 2.1 Sliding window minimums

Algorithm 2 details our proposal for computing all minimal values of a sliding window of a vector of integers *v* in linear time and with zero memory allocation. Note that modifying this algorithm for computing the maximal values instead of minimal values is straightforward. Section 3.1, we propose results showing the advantages of this proposed algorithm.

##### Algorithm 2

sliding minimum window

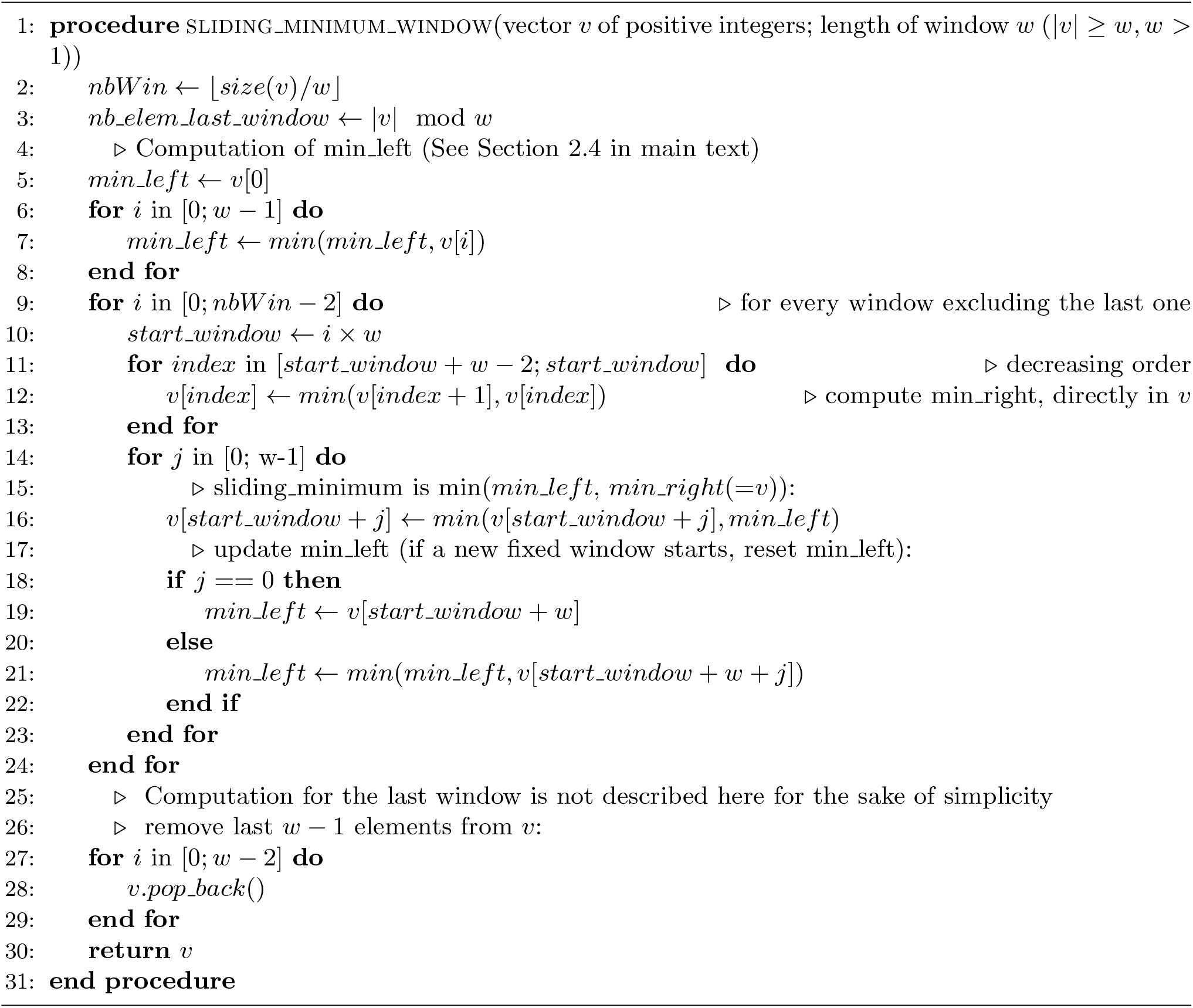

#### 2.2 Skip stretches of consecutive absent *k*-mers

Knowing the absence of a *s*-mer allows deducing that all *k*-mers containing this *s*-mer are absent. This allows inferring the existence of a stretch of consecutive absent *k*-mers.

We exploit this simple idea further. If one detects that two absent *s*-mers are *z* + 1 positions away in the query, then any *k*-mer starting at any position between them is also absent. In the `fimpera` algorithm, if a *s*-mer is not found during the query, an optimization consists of searching for the abundance of the *s*-mer *z* + 1 positions further away in the query. If that *s*-mer is also absent, there is no need to query any *s*-mer in between.

Thus, `fimpera` only needs to query one *s*-mer every *z* +1 position as long as the queried *s*-mers are absent in the counting AMQ, effectively saving time.

We can now propose the entire algorithm of `fimpera` (algorithm 3), using the sliding minimal window algorithm and including the optimization of skipping stretches of absent *k*-mers. The skip optimization occurs line 31: if a negative *s*-mer is called, the algorithm jumps *z* + 1 position away in the sequence, probing for another absent *s*-mer. A positive answer triggers line 17, backtracking *z* positions backward. In this situation, we keep track of the fact that we are skipping *s*-mers via the *extending stretch* flag.

##### Algorithm 3

Optimized `fimpera’s` query (*q* ∈ ∑*; cAMQ indexing *s*-mers; *k* and *z* in ℕ^+^) with |*q*| ≥ *k, z* < *k*.

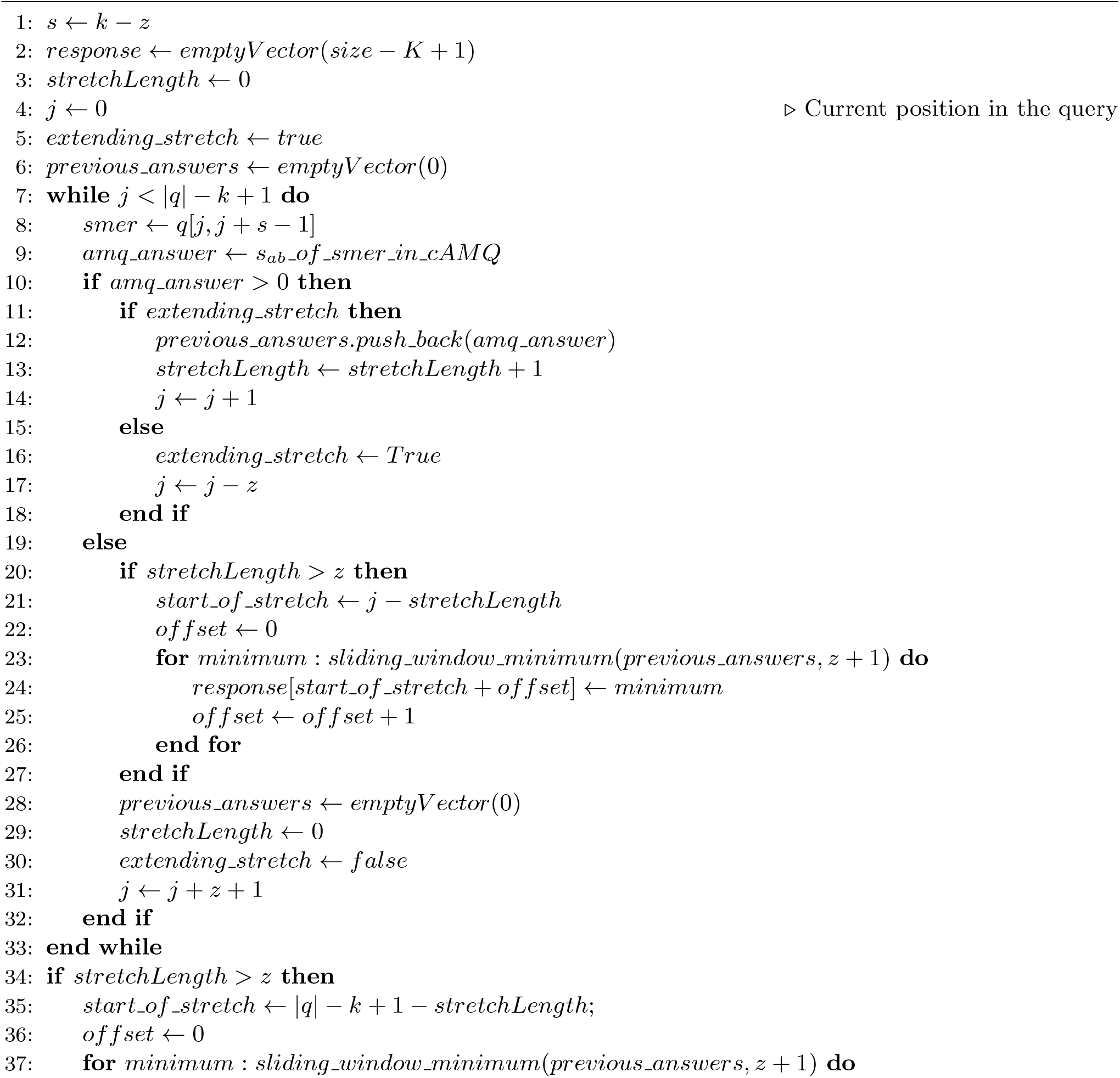
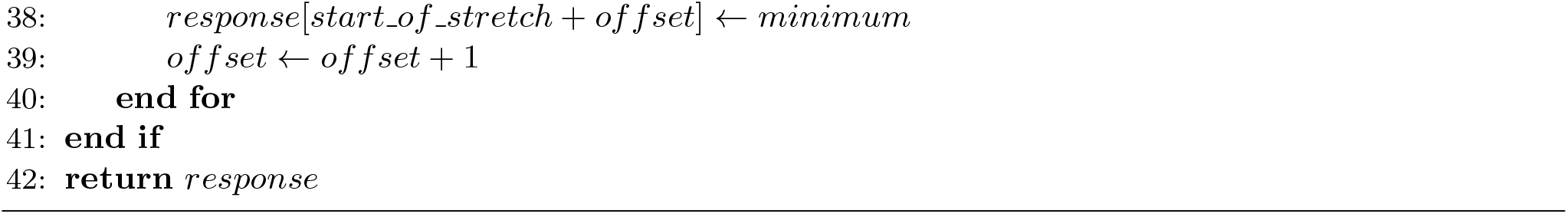

### 3 Additional results

In this section we propose additional results, highlighting the advantages of the new algorithm we propose for computing the minimal value of each sliding window (Section 3.1), or studying the connection between using `fimpera` or using a simple bloom filter with *z* + 1 hash functions (Section 3.2). Finally, we propose additional results focused on the effect of using the *s*_*ab*_ instead of the abundance of *s*-mers (Section 3.3).

#### 3.1 Sliding window minimum benchmark

We propose a benchmark of the following algorithms to compare algorithms that find the minimal value of each sliding window on a vector of integers *v*:

1. “*recomputing*”: a naive algorithm, computing naively the minimum in each window. Time: 𝒪 (*size window* × |*v*|).

2. “*recomputing from last min*”: keeping a reference to the minimum *m* in the previous window and recomputing the minimum in the current window only if the element of value *m* was the first element of the previous sliding window. Simply update *m* if the new element in the current window is smaller than *m*. Time: 𝒪 (*size window* × |*v*|) at worst (if *v* is an increasing sequence).

3. a deque-based approach: the deque contains elements of the input vector. For each element *e*, the back of the deque is removed if it is strictly smaller than *e*. Then *e* is added on the back of the deque and elements that are out of the current window are removed. The minimum of the current window is then the front of the deque. Time: O(|*v*|), but requires time consuming memory allocations.

4. “*fixed windows*”: the approach described in Section 2.4, but allocating new vectors. Time: 𝒪 (|*v*|), but requires time consuming memory heap allocations.

5. Our proposal: “*fixed windows in place*”: based on the fixed approach we also introduced (previous item), but without heap allocation, as described in algorithm 2. Time: O(|*v*|).

We propose various tests, highlighting the behaviors of the presented algorithms in different contexts: *v* is made up of random integers with either a small (size 9) or a large (size 100) window, and *v* is made up of increasing integers. In all cases, our proposal is the fastest algorithm. Recall that the implementation, of independent interest, is available at https://github.com/lrobidou/sliding-minimum-windows along with a recipe to reproduce and extend this benchmark.

As shown in Fig. 1, when *size window* is set to 9, approaches can be divided into two groups: those with heap allocations and those without. The fixed window approach with heap allocations is slower than keeping the last minimum (𝒪 (*size window* × |*v*|) at worst), but once heap allocation is prevented (our proposal), this outperforms any other implementation, including keeping the last minimum by a factor 2. Heap allocations are costly: the differences between approaches in 𝒪 (*size window* × |*v*|) with heap allocation and in O(|*v*|) without them are negligible in practice (as long as |*v*| is small enough, see Fig. 2).

Results presented Figure 2 show that the naive approach in O(*size window*×|*v*|) is significantly slower than other approaches, even if it does not require heap allocation.

**Figure 1.**
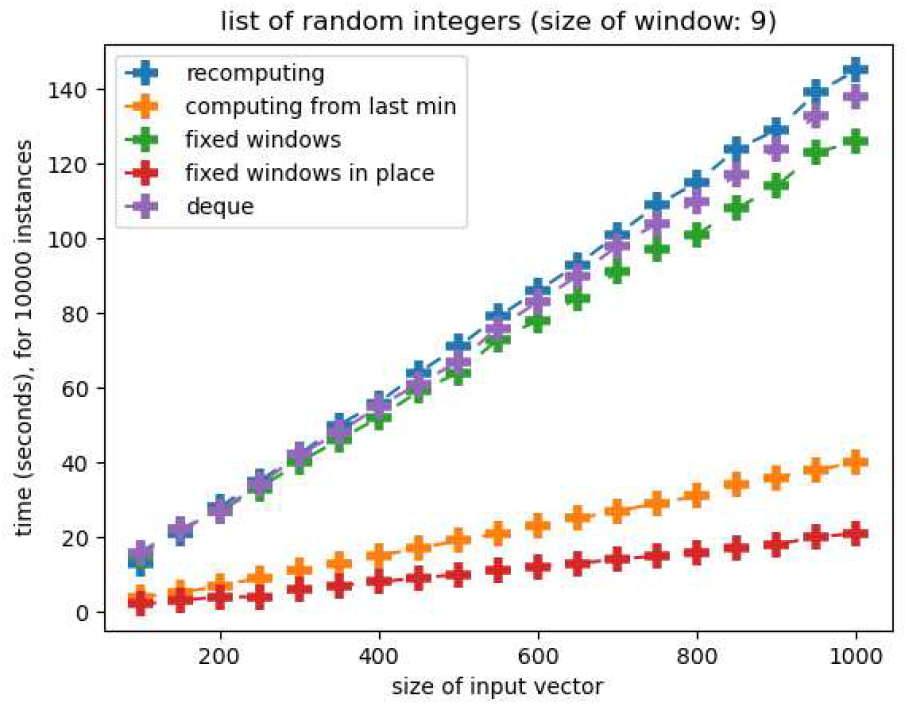
Comparison of five approaches for computing sliding window minimum. The input vector is composed of random integers. The size of the window is set to 9 and for each size of the input vector, 10000 instances were tested.

**Figure 2.**
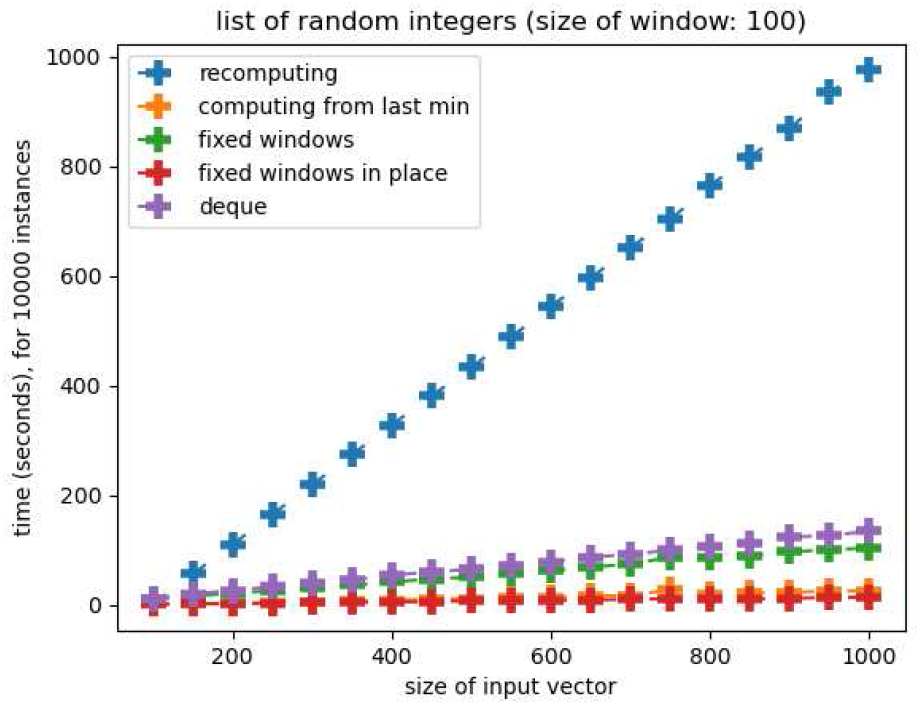
Comparison between 5 approaches for computing sliding window minimum. The input vector is composed of random integers. The size of the window is set to 100.

Keeping a reference to the minimum from the previous window is simpler than using a fixed window but it is two times slower. Moreover, if the input vector contains increasing values, it requires recomputing every window. Indeed, in this case, the minimum from any window is its leftmost element, and as it is out of the next window, the next window needs to be recomputed. See Fig. 3, to see the impact on run time. As illustrated in the figure, the strategy of keeping a reference to the minimum of each window is slower than when applied to random values. In any case, the query time of the method we propose is constant regardless of the input vector’s content.

#### 3.2 Comparing fimpera and a counting Bloom filter with multiple hash functions

As stated in the main manuscript, Section 2.2.1, the `fimpera` strategy may seem similar to associating multiple hash functions to a *k*-mer through its constituent *s*-mers, albeit two consecutive *k*-mers share *z s*-mers, allowing to not query the same *s*-mer multiple times.

In this section, we compared the reported abundance when indexing *k*-mers through *z* + 1 *s*-mers compared to indexing *k*-mers through *z* + 1 independent hash functions.

Using the overestimation score defined in Section 3.4 of the main manuscript, it is possible to compare `fimpera` (*z* = 3) using one hash function (ie. using 4 *s*-mers per *k*-mers) and using 4 independent hash functions. Using the same memory budget, *z* = 3 allows attaining an overestimation score of 98169 while using 4 independent hash functions allows reaching an overestimation score of 57967168 (the lower the better). The false positive rate attained by using *z* = 3 is ≈ 0.36% while using 4 independent hash functions leads to a drastic increase of the false positive rate to 99.94%. This is due to the fact that using 4 independent hash functions saturates the filter and most absent *k*-mers calls lead to false positives.

**Figure 3.**
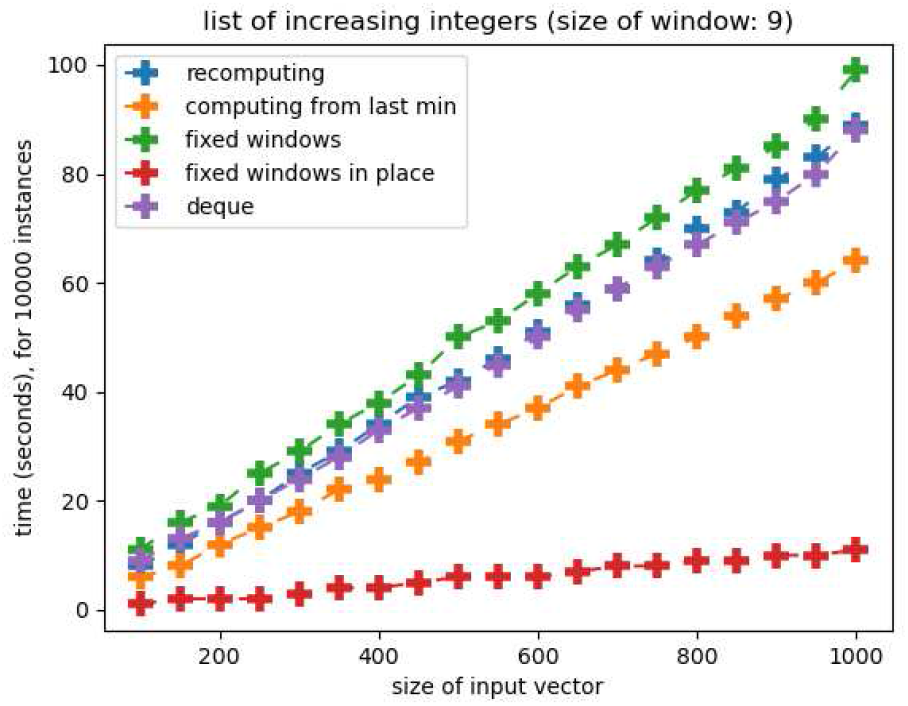
Comparison between 5 approaches for computing sliding window minimum. The size of the window is set to 9 and the input consists of an array of increasing values.

#### 3.3 Comparison between *s*_*ab*_ and abundance of *s*-mers

As stated in Section 2.2.2 of the main manuscript, we index the *s*_*ab*_ of each *s*-mer instead of its abundance in the input dataset. This is motivated by the fact that the *s*_*ab*_ of a *s*-mer α is lower or equal ^1^ to the abundance of α in the input dataset. For instance, consider a *s*-mer α that occurs in two *k*-mers respectively with an abundance of one and two. Then, the abundance of α is three (= 1 + 2), while the *s*_*ab*_ of α is two (= max(1, 2)). Storing the *s*_*ab*_ of α instead of its abundance lowers the abundance overestimations, as it avoids accumulating the abundances of distinct *k*-mers it belongs to.

However, computing the *s*_*ab*_ is time-consuming, as it requires parsing all *s*-mers given an input composed of counted *k*-mers. Thus, keeping compatibility with most *k*-mer counter requires either that the indexation step is performed by `fimpera`, or the use of the abundance of *s*-mers instead of the *s*_*ab*_.

To measure overestimations achieved when indexing the abundance of *s*-mers instead of their *s*_*ab*_, we provide additional results (same datasets as those used in the main manuscript). In the main manuscript, we indexed the llog_2_J values of abundance for limiting the size of the `cBF`. In this section, we show overestimation results computed on exact abundances stored using 8 bits, and thus bounded to 255. We used *k* = 31 and *s* = 28. This is motivated by the fact that, else, no difference could be seen.

In Fig. 4, we show that overestimations introduced by the use of the abundance of each *s*-mer instead of *s*_*ab*_ are limited. This can be numerically estimated:

- the overestimation score comparing a raw counting AMQ to the ground truth is ≈ 8.6 × 10^7^;
- the overestimation score comparing `fimpera` to the ground truth is ≈ 10^4^ (three order of magnitude smaller);
- the additional overestimation score comparing the indexing of the abundance of *s*-mers instead of their *s* − *abundance* with `fimpera` is ≈ 2 × 10^4^ which appears negligible.

**Figure 4.**
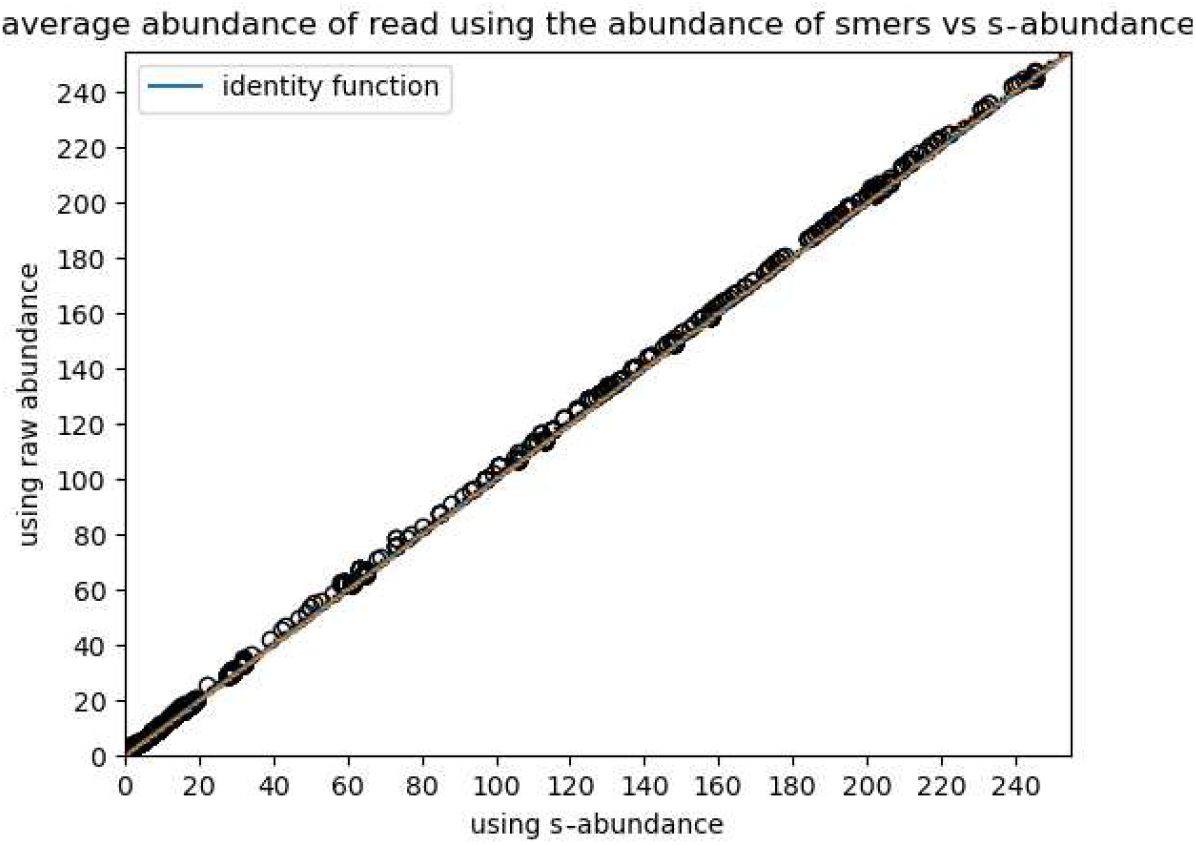
Comparison between the average abundance of *k*-mers from reads query results.

Extremely counter-intuitive case: the abundance of a *s*-mer can be smaller than its *sab*

Actually, in some rare setups, the abundance of *s*-mer can be smaller than its *s*_*ab*_, and even lower than the abundance of any *k*-mer that contains it (which would lead to an underestimation). This is only possible when indexing and querying canonical *k*-mers. This case happens with *s* even, when two *k*-mers having the same canonical form overlap over *s* characters. On a sequence of size *k* + 1 composed of such overlap of two *k*-mers having the same canonical form, their canonical abundance in this sequence is 2, whereas the *s*-mer being the suffix of the first *k*-mer and the prefix of the second has an abundance of 1, thus smaller than its *s*_*ab*_ = *max*(2) = 2.

For instance, consider the sequence *seq* = *TACGTA* and *k* = 5, *s* = 4. The first *k*-mer is *TACGT* and the second *k*-mer is *ACGTA*. Both have the same canonical form *ACGTA*, whose abundance is then equal to two. The *s*-mer they share, *ACGT*, exists only once in *seq*, so its abundance is one, lower than its *s*_*ab*_ equal to two as provided by the abundance of the unique (in this situation) abundance of the canonical *k*-mer it belongs to.

This setup is rare (541 out of 281032928 *s*-mers of the indexed TARA dataset, i.e.: 0.000193 % of its *s*-mers). In practice, this led to zero underestimation of the average of any read. Even if using the abundance of *s*-mers instead of their *s*_*ab*_, and using their canonical versions breaks the theoretical absence of underestimation, we consider the biological impact of these underestimations as perfectly negligible.

#### 3.4 Effects of choosing the *k* value at query time

As shown in the main manuscript in Table 2, the false positive rate of `fimpera` drops with regard to *z* when indexing *k*-mers through their constituent *s*-mers. However it can be useful to index *s*-mers, and then choose *k* at query time. This allows the user to choose any *k* value (≥ *s*) even after the indexation step. The major difference of this approach with respect to having *s* and *k* fixed at indexing time is that, since *k* is unknown when indexing, we cannot compute the s-abundance of *s*-mers. This is due to the fact that computing the s-abundance of *s*-mers requires knowing the *k* value. Thus, in this setup, we rely on the abundance of *s*-mers instead of their s-abundance.

In Table 1, we show that the false positive rate drops with regard to *z*, with a fixed *s* value.

**Table 1:**
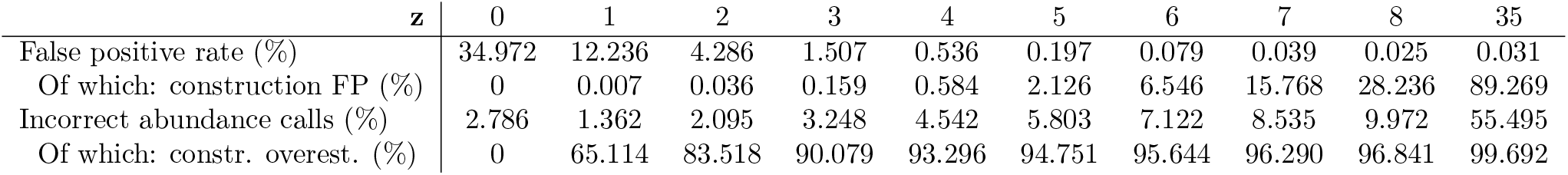
Influence of the *z* parameter on the quality of the results when *s* if fixed. “*constr*.” stands for “*construction*” and “*overest*.” stands for “*overestimation*”. The Incorrect abundance calls are computed over true positive calls only.

As a side note, as previously mentioned, when choosing *k* at query time, one cannot compute the s-abundance of the *s*-mers. Hence, in this setup, the abundance of *s*-mers is higher than the situation when *k* and *z* are known at indexing time, in which one would have used the s-abundance.

Consequently, less *s*-mers have an abundance lower than the threshold and are filtered out, slightly increasing the FP rate, comparatively with results presented in the main manuscript.

Note that choosing *z* at query time allows using a high value, such as 35, while still limiting the construction false positives. This is due to the fact that construction false positive rate mainly depends on the *s* value.

#### 3.5 Effect of grouping values using log functions

In the main manuscript, Section 2.5, we indexed the [log_2_] values of abundance for limiting the size of the `cBF`. In this section, we show overestimation results computed on exact abundances stored using 8 bits, and thus bounded to 255. We used *k* = 31 and *s* = 28.

Fig. 5 shows the response of `fimpera` for each ground truth abundance, not considering abundances as their llog_2_J values. The median of response for each ground truth value is close to that value (most box plots consist in the first quartile, the median and the third quartile being equal to the ground truth). Using the same overestimation score as in Section 3.4 (main manuscript), we compared `fimpera` (*z* = 3) and the underlying counting Bloom Filter. Using the same memory budget, `fimpera` has an overestimation score of 98169 while the counting Bloom Filter has an overestimation score of 85873897 (3 orders of magnitude higher, the lower the better).

**Figure 5.**
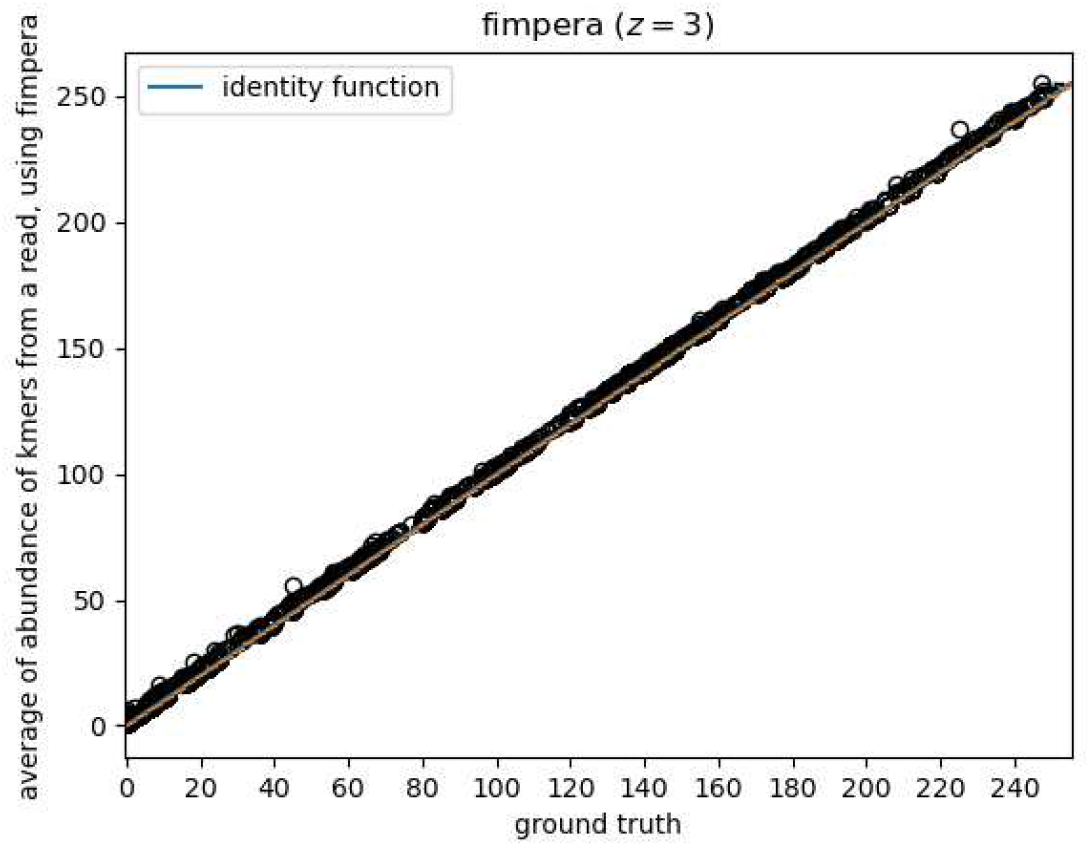
Average abundance of reads reported by `fimpera` (*k* = 31, *s* = 28) using one hash function with regard to the ground truth, compared to the identity function.

Thus, the results of `fimpera` still hold when not using any surjective function to group values, eg. by their log.

## Footnotes

1 except in some extremely rare cases with palindromic *s*-mers as described below

## Notes

https://github.com/lrobidou/fimpera

### Competing Interest Statement

The authors have declared no competing interest.

### Summary of Updates

Addition of two experiments in the supplementary materials Revision of the state of the art section Changes in notations

